# IPCAPS: an R package for iterative pruning to capture population structure

**DOI:** 10.1101/186874

**Authors:** Kridsadakorn Chaichoompu, Fentaw Abegaz Yazew, Sissades Tongsima, Philip James Shaw, Anavaj Sakuntabhai, Luísa Pereira, Kristel Van Steen

## Abstract

**Background:** Resolving population genetic structure is challenging, especially when dealing with closely related or geographically confined populations. Although Principal Component Analysis (PCA)-based methods and genomic variation with single nucleotide polymorphisms (SNPs) are widely used to describe shared genetic ancestry, improvements can be made especially when fine-scale population structure is the target.

**Results:** This work presents an R package called IPCAPS, which uses SNP information for resolving possibly fine-scale population structure. The IPCAPS routines are built on the iterative pruning Principal Component Analysis (ipPCA) framework that systematically assigns individuals to genetically similar subgroups. In each iteration, our tool is able to detect and eliminate outliers, hereby avoiding severe misclassification errors.

**Conclusions:** IPCAPS supports different measurement scales for variables used to identify substructure. Hence, panels of gene expression and methylation data can be accommodated as well. The tool can also be applied in patient sub-phenotyping contexts. IPCAPS is developed in R and is freely available from bio3.giga.ulg.ac.be/ipcaps

## Background

Single Nucleotide Polymorphisms (SNPs) can be used to identify population substructure, but resolving complex substructures remains challenging [1]. Owing to the relatively low information load carried by single SNPs, usually thousands of them are needed to generate sufficient power for effective resolution of population strata due to shared genetic ancestry [2]. Moreover in practice with high-density genome-wide SNP datasets, linkage disequilibrium (LD) and haplotype patterns are likely to exist, which can be exploited for the inference of population structure [3]. On the one hand, exploiting haplotype patterns is potentially informative, but comes with a high computational burden. On the other hand, although removing LD by pruning strategies can eliminate some spurious substructure patterns, it may limit our ability to identify subtle subgroupings.

The identification of substructure in a genome wide association study sample of healthy controls or patients is a clustering problem. Conventional population structure analyses use Bayesian statistics to show relationships amongst individuals in terms of their so-called admixture profiles, where individuals can be clustered by using ratios of ancestral components, see also [4]. The iterative pruning Principal Component Analysis (ipPCA) approach differs from this paradigm as it assigns individuals to subpopulations without making assumptions of population ancestry [5]. At the heart of ipPCA lies performing PCA with genotype data, similar to EIGENSTRAT [2]. If substructure exists in a principal component (PC) space (ascertained using, for instance, Tracy-Widom statistics [5], or the EigenDev heuristic [6]), individuals are assigned into one of two clusters using a 2-means algorithm for which cluster centers are initialized with a fuzzy c-means algorithm. The test for substructure and clustering is performed iteratively on nested datasets until no further substructure is detected, i.e. until a stopping criterion based on fixation index (F_ST_) is satisfied. F_ST_ is commonly used to measure genetic distance between populations. The software developed to perform ipPCA has some shortcomings though. Notably, it is limited to a MATLAB environment, which is not freely available. Also, outliers can severely disturb the clustering analysis. These limitations are addressed in IPCAPS, which improves the power of fine-scale population structure, while appropriately identifying and handling outliers.

## Implementation

The R package IPCAPS provides one synthetic dataset and seven functions:

1. simSNP: a synthetic dataset containing SNPs and population labels.
2. ipcaps: a function for unsupervised clustering to capture population structure based on iterative pruning.
3. rubikClust: a function for unsupervised clustering to detect rough structures and outliers.
4. cal.PC.linear: a function for linear PCA.
5. fst.hudson: a function for average F_ST_ calculation between two groups.
6. fst.each.snp.hudson: a function for F_ST_ calculation for all SNPs between two groups.
7. plot.3views: a function to create scatter plots in three views.
8. top.discriminator: a function to detect top discriminators between two groups.

See the IPCAPS reference manual for details of the functions, arguments, default settings, and for optional user-defined parameters.

The IPCAPS package implements unsupervised strategies that facilitate the detection of fine-scale structure in samples, extracted from informative genetic markers. For general populations, information regarding substructure may come directly from SNPs. For patient samples, general population structure should first be removed via regressing out ancestry informative markers prior to clustering. The latter is incorporated in IPCAPS. Currently, IPCAPS accepts three data input formats: text, PLINK binary (BED, BIM, FAM), and RData (more details in Table 1). In the sequel, we will assume the availability of a sufficiently large SNP panel that is called on a collection of population samples.

**Table 1:** Input formats supported by the function ipcaps.

| Input formats | Descriptions |
| --- | --- |
| PLINK binary format | PLINK binary format consist of 3 files; bed, bim, and fam. To generate these files from PLINK, use option --make-bed |
| Text format | The functions ipcaps supports SNPs in additive coding (0=AA, 1=AB, 2=BB). Each row represents SNP and each column represents individual. SNPs need to be separated by a space or a tab. A big text file should be divided into smaller files to load faster. To input several files, set the option as, for example, files=c('input1.txt','input2.txt','input3.txt') |
| RData format | In case of re-analysis, it is convenient to rerun the function ipcaps using the file rawdata.RData generated by the function ipcaps itself. This file contains a vector of labels and a matrix of SNPs containing N rows of individuals and M columns of SNPs. |

Prior to clustering with IPCAPS, adequate data quality control (QC) steps need to be taken. These are not supported by IPCAPS itself but can easily be performed in PLINK (1.9) [7]. Suggested PLINK parameters include: restrict to founders (–filter-founders), select chromosome 1-22 (–not-chr 0,x,y,xy,mt), perform LD pruning (–indep-pairwise 50 5 0.2), test for Hardy–Weinberg equilibrium (–hwe 0.001), use call rate at least 95% (–mind 0.05), filter out missing SNP above 2% (–geno 0.02), and remove low minimum allele frequency (–maf 0.05). The remaining missing genotype values are SNP-wise imputed by medians.

Rather than performing 2-means clustering in PCA-space, at each iteration, IPCAPS clustering potentially involves the consecutive application of 2 clustering modules. The first, which we call rubikClust, is applied in the 3-dimensional space determined by the 3 first principal components (axes) at an iteration step. It involves applying rotations in 3D by consecutively performing rotations around PC1, PC2, PC3, and may provide >2 clusters. Notably, this approach also allows for a rapid identification of outliers. When samples cannot be divided into 2 groups in this way, the existing R function mixmod (package Rmixmod) is used for latent subgroup detection. In particular, earlier computed PCs (untransformed) at a particular iteration are subjected to multivariate Gaussian mixture modeling and Clustering EM (CEM) estimation [8], allowing for up to three clusters at each iteration.

The iterative loop of IPCAPS can be terminated automatically by calling one of three possible stopping criteria: the number of subgroups is lower than a minimum, the fixation index (F_ST_) is lower than a threshold, and EigenFit is lower than a pre-specified cutoff. The EigenFit criterion is defined by the differences between the logarithms of consecutive eigenvalues, sorted from high to low.

All IPCAPS results are saved in a single directory including textual information about cluster allocations, and visual information such as PC plots and hierarchical trees of group membership. Due to memory restrictions in R, large datasets (i.e., large number of subjects) may need to be split in multiple files and loaded into computer memory via the IPCAPS option files, after which they are internally merged again for iterative PCA. Extra attention is paid on efficient PC calculation [9], also relying on the R package rARPACK.

The analysis procedure using IPCAPS proceeds as follows: Firstly, genotype data are loaded and are analyzed automatically by the function ipcaps. Secondly, cluster membership is returned once clustering process is done. Clusters containing few members are counted as outlying individuals. Lastly, top discriminators between clusters are identified.

Usage example:

\# 1) perform clustering (see Availability of data and materials)

bed.file <- “simSNP.bed” #the bim file and the fam file are required

sample.info <- “simSNP_individuals.txt”

column.number = 2

output.path <- “result”

clusters <- ipcaps(bed=bed.file, label.file=sample.info, lab.col=column.number, out=output.path)

\# 2) Check clustering result

print(clusters$cluster$group)

table(clusters$cluster$label, clusters$cluster$group)

\# 3) Identify top discriminators between groups, for example, group 4 and group 5

bim.file <- “simSNP.bim”

top.snp <-top.discriminator(clusters,4,5,bim.file) head(top.snp)

## Results

We simulated genotype data for 10,000 independent SNPs and 760 individuals belonging to one of three populations (250 individuals each) and 10 outliers (see Availability of data and materials). The pairwise genetic distance between populations was set to F_ST_=0.005 [10]. Ten outlying individuals were generated by replacing the 1st and the 2nd eigenvectors by extreme values, and then the SNP matrix was reconstructed using the singular value decomposition formula [11]. Two-dimensional PC plots of the first 3 PCs only reveals a separation between populations (with overlap) for PC2 versus PC3 (Fig. 1-A). However, application of IPCAPS on the simulated data and thus flexible use of PC information and clustering stopping rules as described before, could clearly identify sample substructure (Fig. 1-B). Non-outlying individuals were correctly assigned to their respective subgroups. In a real-life data application, we considered four populations of HapMap (CEU, YRI, CHB, and JPT) [12]. These populations have been considered before in the evaluation of non-linear PCA to detect fine substructure [13]. After data QC as described before, 132,873 SNPs and 395 individuals remained (see Availability of data and materials). Using classic PCA, visualizing data into two-dimensional space based on the first two PCs is not enough to fully describe substructures. Whereas non-linear PCA is able to provide a hierarchical visualization with only the first 2 PCs, as claimed by the authors [13], including PC3 clearly improves the detection of substructure of four strata, but the authors do not give recommendations on how to select the optimal number of non-linear PCs (Fig. 1-C). The iterative approach adopted in IPCAPS can distinguish populations for which the internal substructure becomes increasingly finer: CEU, YRI, CHB, and JPT populations are well separated by IPCAPS, which also separates the genetically rather similar population CHB and JPT, with only one misclassified subject. In addition, we obtained 560 unique SNPs after combining the top discriminators among four main groups, while outliers were ignored (Fig. 1-D).

**Fig. 1.**
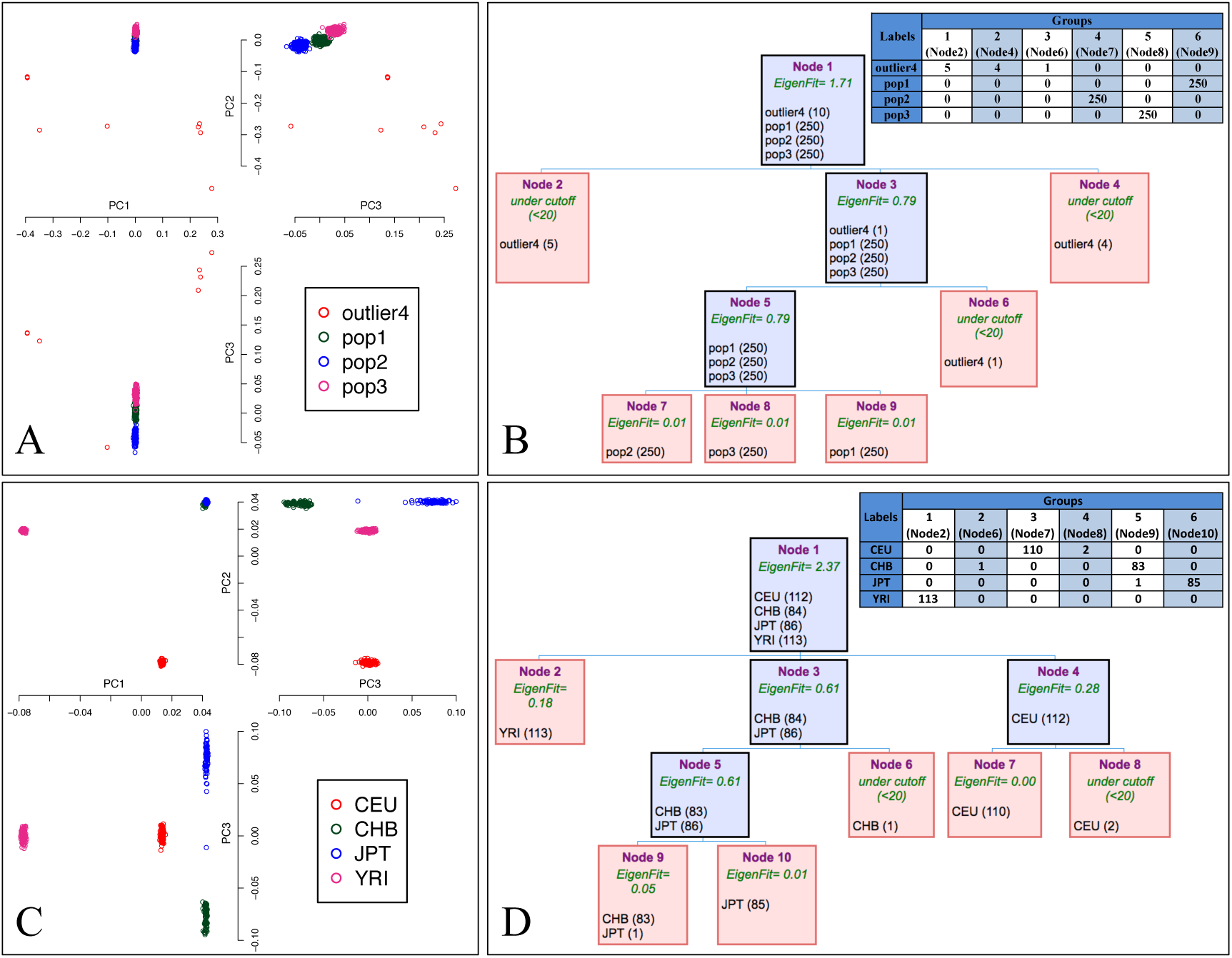
The output from IPCAPs. (A) PC plot of iteration 1 for synthetic data (B) a typical tree output and a summary table for synthetic data (C) PC plot of iteration 1 for the HapMap data (D) a typical tree output and a summary table for the HapMap data. For (B) and (D), the intermediate results are in blue, and the final clusters are in red.

## Conclusions

Fine-scale resolution of population substructure can be captured using independent SNPs once all redundancies are filtered out. In this work, we have introduced a flexible and efficient R package to accomplish an unsupervised clustering without prior knowledge, in the search for strata of individuals with similar genetic profiles. The tool performs well in fine-scale and broad-scale resolution settings.

The IPCAPS routines allow relatively easy extension to input data derived from transcriptome or epigenome experiments.

## Availability and requirements

Project name: IPCAPS

Project home page: bio3.giga.ulg.ac.be/ipcaps

Operating system: Platform independent

Programming language: R version >= 3.0.0

Other requirements: Dependency R packages; RMatrix, expm, fpc, Rmixmod, LPCM, apcluster, rARPACK, igraph

License: GPLv3

## Abbreviations

F_ST_: fixation index
LD: linkage disequilibrium
PC: principal component
PCA: principal component analysis
QC: quality control
SNP: single nucleotide polymorphisms

## Availability of data and materials

The datasets generated during and/or analyzed during the current study are available in the BIO3’s website, bio3.giga.ulg.ac.be/ipcaps

## Authors’ contributions

KC and KVS conceived the methodology. KC designed the software and implemented the R code for the software. FAY, ST and PJS suggested additional features. All authors contributed to write the paper, read, and approved the final manuscript.

## Acknowledgements

The authors thank Pongsakorn Wangkumhang, and Alisa Wilantho for helpful discussions. We also thank Chumpol Ngamphiw, Raphaël Philippart, and Alain Empain for critical help on computing clusters.

## References

1. Neuditschko M, Khatkar MS, Raadsma HW. NetView: a high-definition network-visualization approach to detect fine-scale population structures from genome-wide patterns of variation. PloS One. 2012;7:e48375.

2. Price AL, Patterson NJ, Plenge RM, Weinblatt ME, Shadick NA, Reich D. Principal components analysis corrects for stratification in genome-wide association studies. Nat. Genet. 2006;38:904–9.

3. Lawson DJ, Hellenthal G, Myers S, Falush D. Inference of population structure using dense haplotype data. PLoS Genet. 2012;8:e1002453.

4. Corander J, Marttinen P, Sirén J, Tang J. Enhanced Bayesian modelling in BAPS software for learning genetic structures of populations. BMC Bioinformatics. 2008;9:539.

5. Intarapanich A, Shaw PJ, Assawamakin A, Wangkumhang P, Ngamphiw C, Chaichoompu K, et al. Iterative pruning PCA improves resolution of highly structured populations. BMC Bioinformatics. 2009;10:382.

6. Limpiti T, Intarapanich A, Assawamakin A, Shaw PJ, Wangkumhang P, Piriyapongsa J, et al. Study of large and highly stratified population datasets by combining iterative pruning principal component analysis and structure. BMC Bioinformatics. 2011;12:255.

7. Chang CC, Chow CC, Tellier LC, Vattikuti S, Purcell SM, Lee JJ. Second-generation PLINK: rising to the challenge of larger and richer datasets. GigaScience [Internet]. 2015;4. Available from: doi.org/10.1186%2Fs13742-015-0047-8

8. Lebret R, Iovleff S, Langrognet F, Biernacki C, Celeux G, Govaert G. Rmixmod: TheRPackage of the Model-Based Unsupervised, Supervised, and Semi-Supervised ClassificationMixmodLibrary. J. Stat. Softw. [Internet]. 2015;67. Available from: doi.org/10.18637%2Fjss.v067.i06

9. Clayton D. snpStats: SnpMatrix and XSnpMatrix classes and methods. 2015.

10. Balding DJ, Nichols RA. A method for quantifying differentiation between populations at multi-allelic loci and its implications for investigating identity and paternity. Genetica. 1995;96:3–12.

11. Liu L, Zhang D, Liu H, Arendt C. Robust methods for population stratification in genome wide association studies. BMC Bioinformatics. 2013;14:132.

12. Gibbs RA, Belmont JW, Hardenbol P, Willis TD, Yu F, Yang H, et al. The International HapMap Project. Nature. 2003;426:789–96.

13. Alanis-Lobato G, Cannistraci CV, Eriksson A, Manica A, Ravasi T. Highlighting nonlinear patterns in population genetics datasets. Sci. Rep. 2015;5:8140.

